# SpatialExperiment: infrastructure for spatially resolved transcriptomics data in R using Bioconductor

**DOI:** 10.1101/2021.01.27.428431

**Authors:** Dario Righelli, Lukas M. Weber, Helena L. Crowell, Brenda Pardo, Leonardo Collado-Torres, Shila Ghazanfar, Aaron T. L. Lun, Stephanie C. Hicks, Davide Risso

**Author notes:** Equal contributions (first authors) (Dario Righelli, Lukas M. Weber, Helena L. Crowell). Equal contributions (senior authors) (Stephanie C. Hicks, Davide Risso).

## Abstract

**Summary:** *SpatialExperiment* is a new data infrastructure for storing and accessing spatially resolved transcriptomics data, implemented within the R/Bioconductor framework, which provides advantages of modularity, interoperability, standardized operations, and comprehensive documentation. Here, we demonstrate the structure and user interface with examples from the 10x Genomics Visium and seqFISH platforms, and provide access to example datasets and visualization tools in the *STexampleData*, *TENxVisiumData*, and *ggspavis* packages.

**Availability and Implementation:** The *SpatialExperiment*, *STexampleData*, *TENxVisiumData*, and *ggspavis* packages are available from Bioconductor. The package versions described in this manuscript are available in Bioconductor version 3.15 onwards.

**Supplementary Information:** Supplementary Tables and Figures are available online.

## Introduction

Spatially resolved transcriptomics (ST) refers to a new set of high-throughput technologies, which measure up to transcriptome-wide gene expression along with the spatial coordinates of the measurements. Technological platforms differ in terms of the number of measured genes (from hundreds to full transcriptome) and spatial resolution (from multiple cells per coordinate to approximately single-cell to sub-cellular). Examples of ST platforms include Spatial Transcriptomics [1], 10x Genomics Visium [2], Slide-seq [3], Slide-seqV2 [4], sci-Space [5], seqFISH [6,7], seqFISH+ [8], osmFISH [9], and MERFISH [10–12]. These can be classified into spot-based and molecule-based platforms. Spot-based platforms measure transcriptome-wide gene expression at a series of spatial coordinates (spots) on a tissue slide (Spatial Transcriptomics, 10x Genomics Visium, Slide-seq, Slide-seqV2, and sci-Space), while molecule-based platforms detect large sets of distinct individual messenger RNA (mRNA) molecules *in situ* at up to sub-cellular resolution (seqFISH, seqFISH+, osmFISH, and MERFISH). ST platforms have been applied to investigate spatial patterns of gene expression in a variety of biological systems, including the human brain [13], mouse brain [14], cancer [15,16], and mouse embryogenesis [5,17]. By combining molecular and spatial information, these platforms promise to continue to generate new insights about biological processes that manifest with spatial specificity within tissues.

However, to effectively analyze these data, specialized and robust data infrastructures are required, to facilitate storage, retrieval, subsetting, and interfacing with downstream tools. Here, we describe *SpatialExperiment*, a new data infrastructure developed within the R/Bioconductor framework, which extends the popular *SingleCellExperiment* [18] class for single-cell RNA sequencing (scRNA-seq) data to the spatial context, with observations taking place at the level of spots or molecules instead of cells. Several recent studies have reused or extended existing single-cell infrastructure to store additional spatial information [13,17]. In addition, several comprehensive analysis workflows have been developed using modified single-cell infrastructure adapted for spatial data, including *Seurat* [19], *Giotto* [20], and *Squidpy* [21]. However, while each of these workflows enables powerful analyses, it remains difficult for users to combine elements in a modular way, since each workflow relies on a separate infrastructure. There does not yet exist a common, standardized infrastructure for storing and accessing ST data in R, which would allow users to easily build workflows combining methods and software developed by different groups. A well-designed independent data infrastructure simplifies the work of various users, including developers of downstream analysis methods who can reuse the structure to store inputs and outputs, and analysts who can rely on the structure to connect packages from different developers into analysis pipelines. By working within the Bioconductor framework, we take advantage of long-standing Bioconductor principles of modularity, interoperability, continuous testing, and comprehensive documentation [18,22]. Furthermore, we can ensure compatibility with existing analysis packages designed for the *SingleCellExperiment* structure for single-cell data, providing a robust, flexible, and user-friendly resource for the research community. In addition to the *SpatialExperiment* package, we provide the *STexampleData* and *TENxVisiumData* packages (example datasets) and *ggspavis* package (visualization tools) for use in examples, tutorials, demonstrations, and teaching.

## Results

The *SpatialExperiment* package provides access to the core data infrastructure (referred to as a class), as well as functions to create, modify, and access instances of the class (objects). Objects contain the following components adapted from the existing *SingleCellExperiment* class: (i) assays, tables of measurement values such as raw and transformed transcript counts (note that within the Bioconductor framework, rows usually correspond to features, and columns to observations); (ii) rowData, additional information (metadata) describing the features (e.g. gene IDs and names); (iii) colData, metadata describing the observations (e.g. barcode IDs or cell IDs); and (iv) reducedDims, reduced dimension representations (e.g. principal component analysis) of the measurements. *SpatialExperiment* objects also contain the following further components to store spatial information: (v) additional metadata stored in colData describing spatial characteristics of the spatial coordinates (spots) or cells (e.g. indicators for whether spots are located within the region overlapping with tissue); (vi) spatialCoords, spatial coordinates associated with each observation (e.g. x and y coordinates on the tissue slide); and, (vii) imgData, image files (e.g. histology images) and information related to the images (e.g. resolution in pixels) (**Figure 1**).

**Figure 1.**
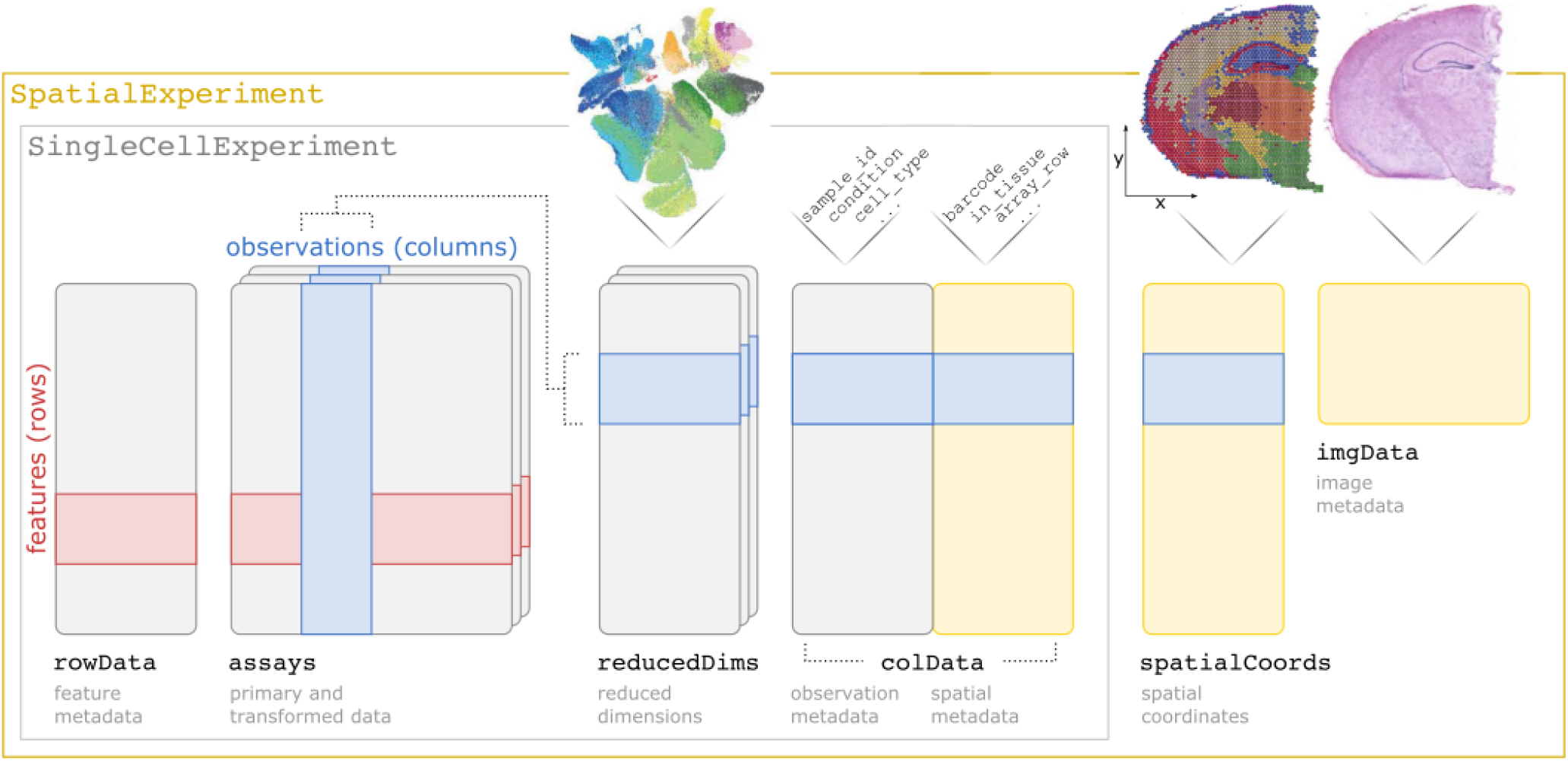
Overview of the *SpatialExperiment* class structure, including assays (tables of measurement values), rowData (metadata describing features), reducedDims (reduced dimension representations), colData (nonspatial and spatial metadata describing observations), spatialCoords (spatial coordinates associated with the observations), and imgData (image files and information).

Accessor and replacement functions allow each of these components to be extracted or modified. Since *SpatialExperiment* extends *SingleCellExperiment*, methods developed for single-cell analyses [18] (e.g. preprocessing and normalization methods from scater [23], downstream methods from scran [24], and visualization tools from iSEE [25]) can be applied to *SpatialExperiment* objects, treating spots as single cells. Spatial coordinates are stored in spatialCoords as a numeric matrix, allowing these to be provided easily to downstream spatial analysis packages in R outside Bioconductor (e.g. packages from geostatistics such as sp [26] and sf [27]), consistent with reducedDims in *SingleCellExperiment*. For spot-based data, assays contains a table named counts containing the gene counts, while for molecule-based data, assays may contain two tables named counts and molecules containing total gene counts per cell as well as molecule-level information such as spatial coordinates per molecule (formatted as a BumpyMatrix [28]). For datasets that are too large to store in-memory, *SpatialExperiment* can reuse existing Bioconductor infrastructure for sparse matrices and on-disk data representations through the *DelayedArray* framework [29]. Image information is stored in imgData as a table containing sample IDs, image IDs, any other information such as scaling factors, and the underlying image data. The image data can be stored as either a fully realized in-memory object (for small images), a path to a local file that is loaded into memory on demand (for large images), or a URL to a remotely hosted image that is retrieved on demand. *SpatialExperiment* objects can be created with a general constructor function, SpatialExperiment(), or alternatively with a dedicated constructor function for the 10x Genomics Visium platform, read10xVisium(), which creates an object from the raw input files from the 10x Genomics Visium Space Ranger software [30]. For Visium data, colData includes the columns in_tissue, array_row, and array_col. Measurements from multiple biological samples can be stored within a single object, and linked across the components by providing unique sample IDs. In addition, we provide the associated data packages *STexampleData* (example datasets from several platforms) and *TENxVisiumData* (publicly available Visium datasets provided by 10x Genomics), and the *ggspavis* package providing visualization functions designed for *SpatialExperiment* objects (**Supplementary Figure 1** and **Supplementary Table 1**).

## Discussion

Standardized data infrastructure for scRNA-seq data (e.g. *SingleCellExperiment* [18] within the R/Bioconductor framework) has greatly streamlined the work of data analysts and downstream method developers. For example, relying on common formats for inputs and outputs from individual packages allows users to connect packages into complete analysis pipelines, and operations such as subsetting by row (gene) or column (barcode or cell) across the entire object helps avoid errors. For single-cell data, this has enabled the development of comprehensive workflows and tutorials [18,31], which are an invaluable resource for new users. Here, we provide a new data infrastructure for ST data, extending the existing *SingleCellExperiment* class within the Bioconductor framework. In addition, we provide associated packages containing example datasets (*STexampleData* and *TENxVisiumData*) and visualization functions (*ggspavis*), for use in examples, tutorials, demonstrations, and teaching.

Existing alternative infrastructure for ST data includes object classes provided in the *Seurat* [19] and *Giotto* [20] packages in R, and *Squidpy/AnnData* [21,32] in Python, which provide similar underlying functionality such as storing annotated tables of measurement values and related spatial and image information. Compared to these alternatives, a key advantage of *SpatialExperiment* is that it has been developed independently of any individual analysis workflow and is compatible with any downstream analysis packages that use the *SpatialExperiment* or *SingleCellExperiment* class within Bioconductor. This allows analysts to easily build customized, modular workflows consisting of packages developed by various research groups, including the latest methods (which may not yet have been integrated into published workflows) as well as any of the wide variety of methods for single-cell data that have been released through Bioconductor.

ST technologies are still in their infancy, and the coming years are likely to see ongoing development of existing platforms as well as the emergence of novel experimental approaches. *SpatialExperiment* is ideally positioned to be extended to accommodate data from new platforms in the future, e.g. through extensions of the more general underlying *SummarizedExperiment* [33] or by integrating with *MultiAssayExperiment* [34] to store measurements from further assay types (transcriptomics, proteomics or spatial immunofluorescence, or epigenomics) or multiple assays from the same spatial coordinates. For example, the *SingleCellMultiModal* package [35] stores *MultiAssayExperiment* objects containing scRNA-seq and ST data as *SingleCellExperiment* and *SpatialExperiment* objects, respectively. Three-dimensional spatial data [36] or data from multiple timepoints could be accommodated within *SpatialExperiment* by storing additional spatial or temporal coordinates. Datasets that are too large to store in-memory can be stored using existing Bioconductor infrastructure for sparse matrices and on-disk data representations through the *DelayedArray* framework [29]. The ability to store image files within the objects (in-memory, locally, or remotely) will assist with correctly keeping track of images in datasets with large numbers of samples, e.g. from consortium efforts. Interoperability between *SpatialExperiment* and other data formats (e.g. in Python) can also be ensured through the use of existing conversion packages such as *zellkonverter* [37] *and LoomExperiment* [38]. *SpatialExperiment* provides the research community with a robust, flexible, and extendable core data infrastructure for ST data, assisting both method developers and analysts to generate reliable and reproducible biological insights from these platforms.

## Acknowledgments

We thank the participants of the EuroBioc2020 “Birds of a Feather” session (14 December 2020) and workshop (16 December 2020) on the topic of infrastructure for ST data in Bioconductor, as well as the members of the *spatial* and *SpatialExperiment* channels of the Bioconductor community Slack workspace, for helpful feedback and suggestions.

## Author contributions

D. Righelli, LMW, and HLC designed the *SpatialExperiment* class structure, with input from all other authors. D. Righelli led the implementation of the *SpatialExperiment* class, with significant code input from HLC. LMW developed the example data package *STexampleData* and the visualization package *ggspavis*. HLC developed the data package *TENxVisiumData* and provided functions for the *ggspavis* package. BP and LCT tested an earlier version of the *SpatialExperiment* class and provided input on design choices for the final class structure. SG provided input and examples for applying the *SpatialExperiment* class to molecule-based ST data. ATLL provided input on design choices for the *SpatialExperiment* class structure. SCH and D. Risso provided supervision and input on design choices for the *SpatialExperiment* class structure. LMW drafted the paper with input from all other authors. All authors approved the final version of the manuscript.

## Code and data availability

The *SpatialExperiment* package is available from Bioconductor at https://bioconductor.org/packages/SpatialExperiment. The *STexampleData*, *TENxVisiumData*, and *ggspavis* packages are available from Bioconductor at https://bioconductor.org/packages/STexampleData, https://bioconductor.org/packages/TENxVisiumData, and https://bioconductor.org/packages/ggspavis respectively. The package versions described in this manuscript are available in Bioconductor version 3.15 onwards. Installation instructions for the release and development versions are also provided on GitHub at https://github.com/drighelli/SpatialExperiment. Datasets from Supplementary Tables 1 and 2 and Supplementary Figure 1 are available as *SpatialExperiment* objects from the *STexampleData* and *TENxVisiumData* packages, and the full original datasets are available from the sources listed in Supplementary Tables 1 and 2 [13,17,39,40].

## Conflicts of interest

The authors declare that they have no financial conflicts of interest.

## Funding

This work was supported by CZF2019-002443 (LMW, D. Righelli, SCH, D. Risso) from the Chan Zuckerberg Initiative DAF, an advised fund of Silicon Valley Community Foundation. LMW, SCH and LC-T were supported by NIH/NIMH U01MH122849 to SCH and LC-T. D. Risso was supported by “Programma per Giovani Ricercatori Rita Levi Montalcini” granted by the Italian Ministry of Education, University, and Research and by the National Cancer Institute of the National Institutes of Health (2U24CA180996). SG was supported by a Royal Society Newton International Fellowship (NIF\R1\181950).

## Supplementary Tables

**Supplementary Table 1.** Summary of example datasets provided in *SpatialExperiment* format in the *STexampleData* package. Table columns describe characteristics for each dataset, and provide the original references. For the *Visium_humanDLPFC* and *seqFISH_mouseEmbryo* datasets, the objects in the *STexampleData* package contain small subsets of the full original datasets, allowing users to easily download and load these datasets for examples and tutorials. The full datasets can be obtained from the original references.

| Dataset name | Platform | Type | Tissue | Number of samples | Number of spots or cells | Number of features (genes) | Contains ground truth labels? | Contains image data? | Source |
| --- | --- | --- | --- | --- | --- | --- | --- | --- | --- |
| Visium_humanDLPFC | 10x Genomics Visium [2] | Spot-based | Human brain | 1 | 3,639 | 33,538 | Yes | Yes | [13,39] |
| Visium_mouseCoronal | 10x Genomics Visium [2] | Spot-based | Mouse brain | 1 | 2,702 | 32,285 | Yes | Yes | [40] |
| seqFISH_mouseEmbryo | seqFISH [6,7] | Molecule-based | Mouse embryo | 1 | 11,026 | 351 | No | No | [17] |
| ST_mouseOB | Spatial Transcriptomics [1] | Spot-based | Mouse brain | 1 | 262 | 15,928 | Yes | No | [1] |
| SlideSeqV2_mouseHPC | Slide-seqV2 [4] | Spot-based | Mouse brain | 1 | 53,208 | 23,264 | Yes | No | [4,41] |

**Supplementary Table 2.** Summary of example datasets provided in *SpatialExperiment* format in the *TENxVisiumData* package. All data are spot-based, and were obtained using the 10x Genomics Visium platform [2]. Table columns describe characteristics for each dataset. For some datasets, targeted expression panels were measured in addition to whole-transcriptome analysis; these are indicated with the name of the panel and corresponding number of genes in italics. The original datasets can be obtained from [42].

| Dataset name | Tissue | Number of samples | Targeted panel(s) | Number of spots | Number of genes |
| --- | --- | --- | --- | --- | --- |
| HumanBreastCancerIDC | Human invasive ductal carcinoma breast | 2 | – | 7,785 | 36,601 |
| HumanBreastCancerILC | Human invasive lobular carcinoma breast | 1 | –<br>*Immunology* | 4,325 | 36,601<br>*1,056* |
| HumanCerebellum | Human cerebellum | 1 | –<br>*Neuroscience* | 4,992 | 36,601<br>*1,186* |
| HumanColorectalCancer | Human invasive adenocarcinoma of the large intestine | 1 | –<br>*Gene signature* | 3,138 | 36,601<br>*1,142* |
| HumanGlioblastoma | Human glioblastoma multiforme | 1 | –<br>*Pan-cancer* | 3,468 | 36,601<br>*1,253* |
| HumanHeart | Human heart | 1 | – | 4,247 | 36,601 |
| HumanLymphNode | Human lymph node | 1 | – | 4,035 | 36,601 |
| HumanOvarianCancer | Human ovarian endometrial adenocarcinoma | 1 | –<br>*Immunology*<br>*Pan-cancer* | 3,493 | 36,601<br>*1,056*<br>*1,253* |
| HumanSpinalCord | Human spinal cord | 1 | –<br>*Neuroscience* | 2,812 | 36,601<br>*1,186* |
| MouseBrainCoronal | Mouse brain (coronal plane) | 1 | – | 2,702 | 32,285 |
| MouseBrainSagittalAnterior | Mouse brain (sagittal slice of the posterior) | 2 | – | 5,520 | 32,285 |
| MouseBrainSagittalPosterior | Mouse brain (sagittal slice of the anterior) | 2 | – | 6,644 | 32,285 |
| MouseKidneyCoronal | Mouse kidney | 1 | – | 1,438 | 32,285 |

## Supplementary Figures

**Supplementary Figure 1.**
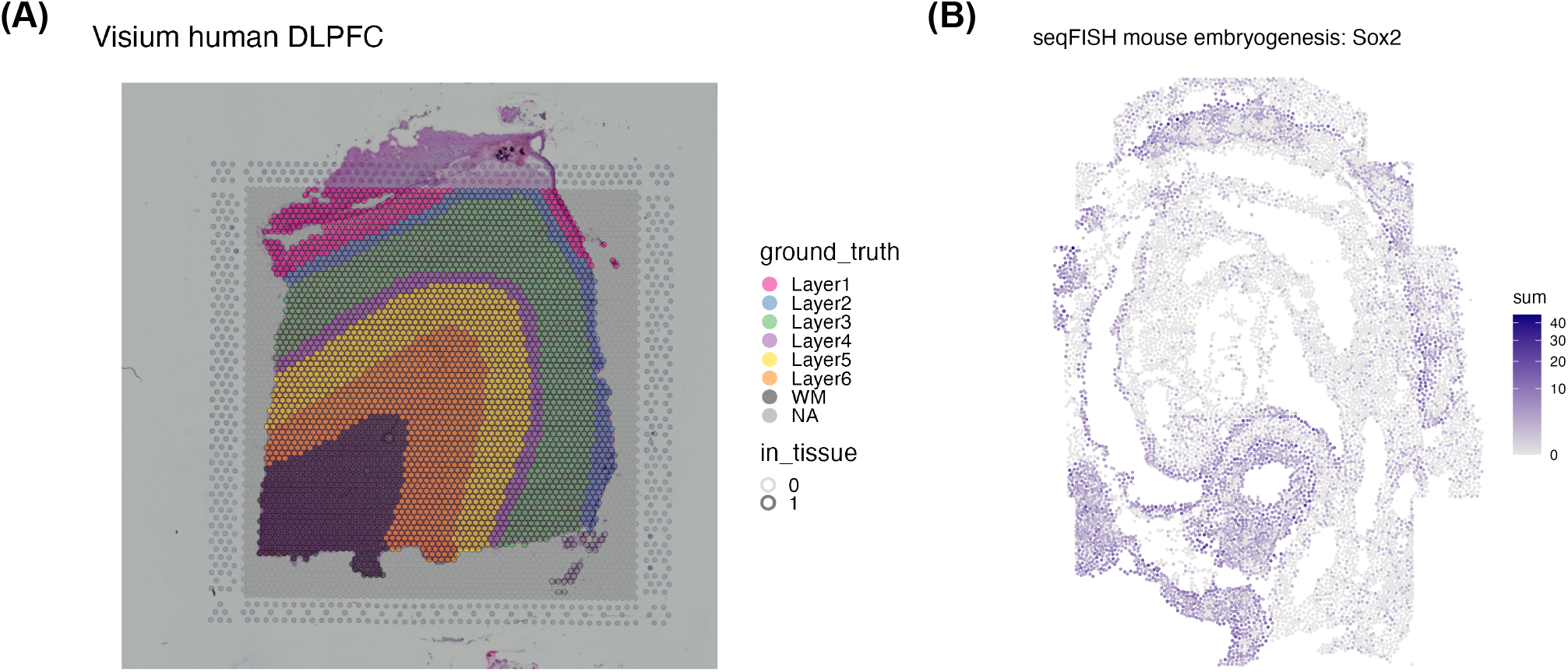
**(A)** Example of visualization of spot-based ST data (*Visium_humanDLPFC* object from the *STexampleData* package). Image shows a histology image as background, grid of spatial coordinates (spots), highlighting for spots that overlap with tissue, and colors for ground truth cluster labels. The dataset represents a single biological sample (sample 151673) from the human brain dorsolateral prefrontal cortex (DLPFC) region [13,39], measured with the 10x Genomics Visium platform. The full dataset contains 12 biological samples, and is available in *SpatialExperiment* format in the *spatialLIBD* Bioconductor package [13,39]. **(B)** Example of visualization of molecule-based ST data (*seqFISH_mouseEmbryo* object from the *STexampleData* package). Color scale shows total mRNA counts per cell for the *Sox2* gene. The dataset represents a subset of cells (embryo 1, z-slice 2) from a published dataset investigating mouse embryogenesis [17], generated using the seqFISH platform. Additional details on the datasets are provided in Supplementary Table 1. Figures were generated using plotting functions from the *ggspavis* package.

## Notes

### Competing Interest Statement

The authors have declared no competing interest.

